# Assessing reliability and accuracy of qPCR, dPCR and ddPCR for estimating mtDNA copy number in songbird blood and sperm cells

**DOI:** 10.1101/2024.11.08.622696

**Authors:** Laima Bagdonaitė, Erica H. Leder, Jan T. Lifjeld, Arild Johnsen, Quentin Mauvisseau

## Abstract

Mitochondrial DNA copy number varies across species, individuals and cell types. Two avian cell types carrying a relatively low number of mitochondria are the red blood cells and spermatozoa. While previous studies investigating variation of mitochondrial abundance in animal sperm have generally used quantitative PCR (qPCR), this method shows potential limitations when quantifying low abundant targets. To mitigate such issues, we investigated and compared the reliability and accuracy of qPCR, digital PCR (dPCR) and droplet digital PCR (ddPCR) to quantify high and low concentration DNA. Using synthetic DNA, we found that both dPCR and ddPCR displayed lower Limit of Detection and Limit of Quantification than qPCR. Using DNA extracted from blood and sperm cells of Eurasian Siskin, we found that qPCR, dPCR and ddPCR reliably quantified mitochondrial DNA in sperm samples, but showed significant differences when analyzing typically lower levels of mtDNA in blood. We found that ddPCR consistently showed lower variation among replicates. These analyses provide critical insights and recommendations for future studies aiming to quantify target mtDNA. Our study indicates that dPCR and ddPCR are the preferred methods when working with samples with low abundance of mtDNA.

## Introduction

Found in most eukaryotic cell types, mitochondria are double-membraned organelles commonly referred to as the “powerhouse of the cell” [1]. The interactions between mitochondrial and nuclear genomes enable oxidative phosphorylation (OXPHOS) processes via which adenosine triphosphate (ATP) is produced, providing energy supporting various cell processes [2–4]. Due to variation in metabolic needs, mitochondria numbers vary tremendously between species, individuals and different cell types [5–7]. For example, the brain and different muscle tissues (often associated with higher metabolic needs), systematically show higher mtDNA copy number than other tissues or cells, such as red blood cells, which typically lack mitochondria in most mammals and are present only in small amounts in birds [3, 6, 8]. Likewise, sperm cells are known to have a much lower mitochondria content than somatic cells [9].

In the animal kingdom, sperm cells consist of three key components: the head, midpiece and tail, the midpiece harboring the mitochondria [10]. Interestingly, bird spermatozoa have been found to contain relatively low amounts of mitochondria per cell, with numbers ranging from 20 to 350 mitochondria, and display large inter and intra specific variation [10]. Passerine birds especially are known for their tremendous variation in sperm morphology, including differences in the shape and volume of different sperm components [11]. While previous interest has been cast on the relationship of sperm midpiece morphology and various sperm properties, such as ATP production, swimming speed or sperm competition [12–15], there remains a scarcity of studies focused on quantifying mitochondrial DNA content in passerine birds such as in [16]. Several studies previously investigated mitochondrial abundance in animal sperm, especially in *Drosophila melanogaster* [17], Chinook salmon [18], humans [19], bull [20] and zebra finch [16], mainly using quantitative polymerase chain reaction (qPCR) techniques.

Quantitative PCR monitors the amplification of the targeted molecule in real time through fluorescence scanning at the end of each amplification cycle, and through the additional analysis of a calibration curve of samples with a known concentration, quantifies the amplicon of interest [21]. A wide range of studies have used qPCR to answer various research questions due to its speed, relative sensitivity and availability [e.g. 22–24]. However, qPCR shows limitations due to the necessity of generating calibration curves to quantify samples, and has been associated with larger variability when analyzing samples at low concentration [25].

Recently, there has been an increase in the utilization of absolute quantification methods, also known as digital PCR (dPCR) and droplet digital PCR (ddPCR). Both methods rely on the partitioning of an initial reaction mixture into thousands of independent droplets, each becoming a separate PCR reaction, in such a way that partitions will contain one or a few target molecules or no target at all [26–29]. One major difference between the partitioning of the sample in the two methods is the use of microfluidic techniques such as micro well plates in dPCR, and the use of oil and emulsification in ddPCR [27, 29]. For both dPCR and ddPCR, following end point PCR, fluorescence signals of each partition are measured to ensure the presence or absence of the targeted amplified DNA molecule. Then, using Poisson statistics, the ratio of positive to the total number of partitions is used to calculate the absolute target concentration (copies/µL) [28, 29]. The last few years have seen commercialization of dPCR and ddPCR technologies with an increase in scientific studies based on using these methods for quantification of target nucleic acids [30, 31] and one of these methods has also been applied to mtDNA in spermatozoa [19]. While digital PCR methods have been argued to be more reproducible than qPCR [32], to our knowledge, no study has investigated the trade-offs between qPCR, dPCR and ddPCR to quantify target mtDNA molecules at low and high concentration.

In this study, we adopt a comparative approach, examining one conventional qPCR method and two absolute quantification methods, dPCR and ddPCR. We aim to evaluate the sensitivity and accuracy of quantifying target mtDNA (Fig. 1) across methods, to identify their quality of results (see four potential scenarios in Fig. 1). For doing so, we used a synthetic mtDNA fragment to investigate the Limit of Detection (LOD) and Limit of Quantification (LOQ) of each method, and additionally compared the mtDNA quantification results and associated variation across replicates in two distinct types of mitochondria-carrying avian biological samples: blood and sperm, associated respectively with low and moderately higher mtDNA numbers.

**Figure 1.**
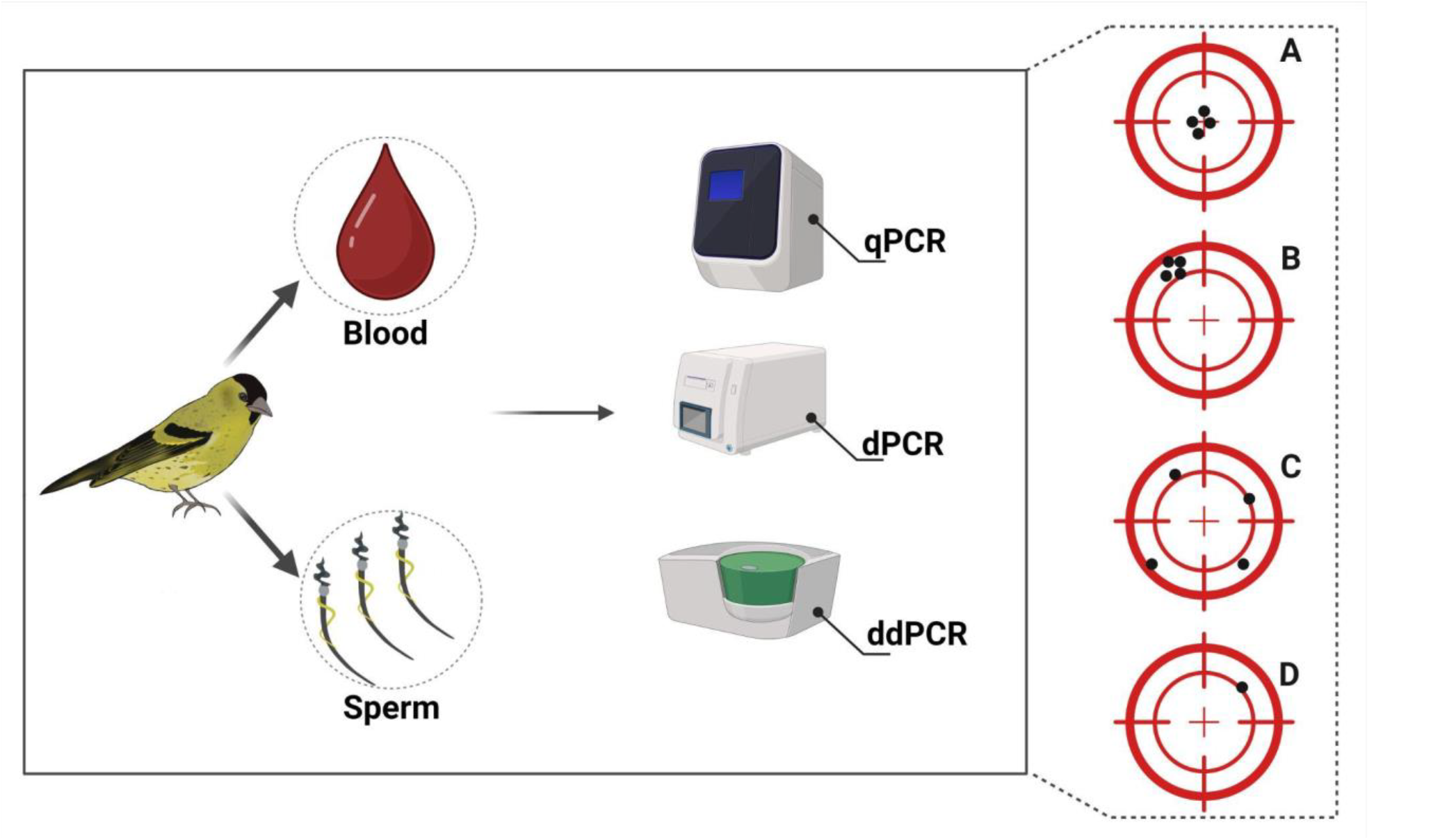
Graphical illustration of the study design. Eurasian siskin (*Spinus spinus*) was selected as the model organism for the study. DNA was extracted from 10 blood and 10 sperm samples of different male individuals. Each sample was quantified in triplicate across all three PCR platforms. The panel on the right (A-D) indicates what level of reliability can be expected when running such assays. A: an optimal scenario with accurate and repeatable target quantification across replicates; B: target quantification is not accurate, but is repeatable across replicates; C: target quantification is not accurate, nor repeatable across replicates; D: target DNA is amplified in only one replicate - target detection is stochastic across replicates and quantification is incorrect, leading to unreliable results. Figure created using BioRender.

## Materials and methods

### Samples

We selected the Eurasian siskin (*Spinus spinus*) as our study species because they are common breeding birds in Norway and are easily attracted to mist nets. Siskins have relatively long sperm cells (219 µm), carrying long midpieces (190 µm) [33], which could indicate they are assembled from a high number of mitochondria [16]. Blood and sperm samples used in this study were collected from male birds during the breeding seasons 2021-2022. A total of 10 blood samples and 10 sperm samples were collected from 5 individuals in 2021 and 13 in 2022 (Supplementary table 1). Birds were caught using mist nets at their breeding sites in southern Norway. Sperm samples were collected via cloacal massage in a similar way as described in Wolfson, 1952. Two males were sampled twice. The ejaculate was collected in a micro capillary tube and immediately mixed with phosphate-buffered saline (PBS) [35]. This was done to prevent the sperm cells from clumping together and to enable pipetting. Sperm samples were then purified through a series of centrifugation and washing with PBS, and frozen at -80°C until DNA extraction. Blood samples were collected through brachial venipuncture (10-20 µL) and stored in 96 % ethanol. Sampling was conducted in adherence to ethical guidelines for use of animals in research and with permission from all relevant local authorities, and approved by the Norwegian Food Safety Authority (permit no. 23294 and 29575), and The Norwegian Environment Agency (permit no. 2021/39021).

**Table 1.**
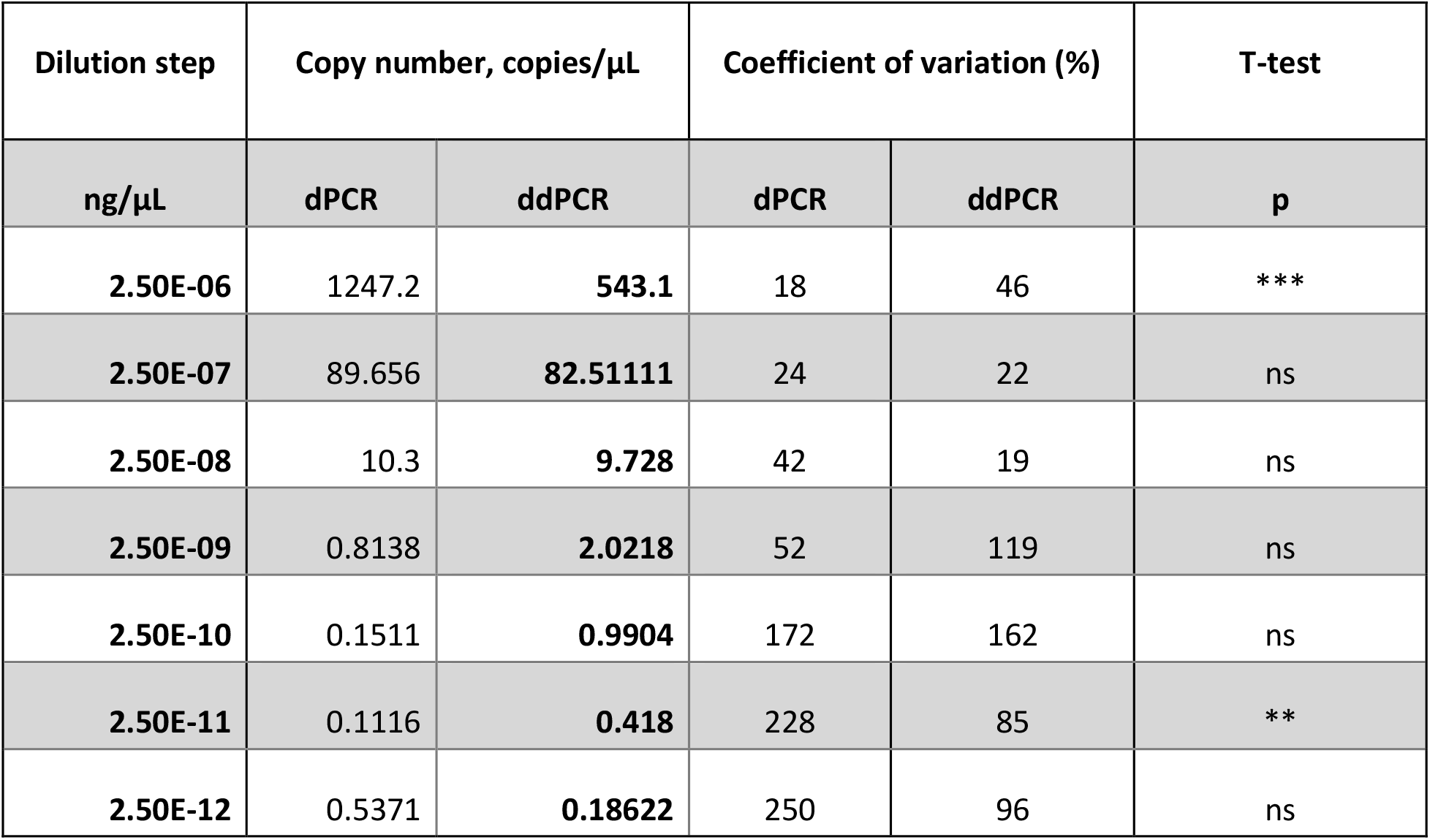
Comparison of synthetic DNA copy number measured with dPCR and ddPCR and the resulting coefficients of variation. The two highest dilution points were excluded due to an excessive number of positive partitions. Variance ratio test (f-test) was first performed to assess which t-test to use. For concentrations at which the variance was equal in both methods (2.5E-06 and 2.5E-11 ng/µL), Student’s t-test was carried out. At all the other concentrations, the Welch t-test was used. Here we provide the results of t-test which compares the mean of concentrations obtained with both methods (“ns” non-significant; ** p-values ≤ 0.01; *** p-values ≤ 0.001).

DNA was extracted from blood samples using the E. Z. N. A. Tissue DNA kit (Omega Bio-Tek, Inc, Norcross, GA, USA) following the manufacturer’s guidelines. DNA was extracted from sperm samples using the QIAamp DNA Micro Kit (QIAGEN, Inc. Valencia, CA, USA) following a protocol for sperm samples as described by Kucera and Heidinger, 2018. We made two changes from the protocol; (i) the mixture of the sample and dithiothreitol (DTT) buffer was incubated for 2 hours in order to ensure the proper lysis of sperm cells and the release of DNA and (ii) the final elution was carried out with 25µL of ddH2O and repeated twice to ensure a maximum DNA yield.

Following DNA extraction, total DNA concentration from each sample was measured with Qubit 2.0 fluorometer with the DNA High Sensitivity Assay (Invitrogen). Sample concentrations were then normalized to a concentration of 0.5ng/µL using ddH2O before further downstream analysis. All DNA extractions were performed in a pre-PCR dedicated laboratory, all the bench surfaces were wiped with 5% deconex solution and absolute ethanol before and after the procedures.

### Primers and synthetic oligo design

Passerine primers (Forward 5’-ATTATTGAGCGAACCCGTCTC-3’ and Reverse 5’-TTCACAGGCAACCAGCTATC-3’) targeting a 98 bp fragment of the mitochondrial 16S gene (5’-ATTATTGAGCGAACCCGTCTCTGTGGCAAAAGAGTGGGATGACTTGTTAGTAGTGGTGAAAAGCCAATCGAGCTGGGTGATAGCTGGTTGCCTGTGAA-3’) were designed following a method similar to that described in [37]. In brief, multiple mitochondrial 16S sequences from a range of songbird species (order Passerides) covering several genera (*Spinus spinus, Emberiza schoeniclus, Coccothraustes coccothraustes, Fringilla coelebs, Pyrrhula pyrrhula, Parus major, Luscinia svecica*) were downloaded from GenBank (http://www.ncbi.nlm.nih.gov) (Table S2). These sequences were then aligned in the Geneious Pro R10 software (https://www.geneious.com) with the multiple alignment function and consensus sequences for each species were generated. Then, the consensus sequences were aligned and primers were created using the primers design function. The specificity of the newly designed primers was visually assessed by inspecting the multi-species alignment and confirmed using the Integrated DNA Technologies (IDT) PrimerQuest tool (PrimerQuest™ program, IDT, Coralville, Iowa, USA). Additional validation was conducted using the primer blast function from the NCBI website (https://www.ncbi.nlm.nih.gov/tools/primer-blast/). Following these *in-silico* validation steps, we performed an additional *in-vitro* validation with PCR and ddPCR on DNA extracted from our target species and other songbird species: *Turdus pilaris, Phylloscopus trochilus, Luscinia svecica, Fringilla montifringilla, Pyrrhula pyrrhula, Regulus regulus, Coccothraustes coccothraustes, Emberiza schoeniclus* (sample information provided in Table S3). These thorough validation steps, including temperature gradient analysis, were done to ensure the specificity and optimal amplification of the target fragment.

### qPCR

Samples were analyzed on a Bio-Rad CFX96 Real-Time System (Bio-Rad Laboratories, California, United States) using the primers described earlier. Quantitative PCR reactions were conducted in a 20 µL final volume containing 10µL QIAcuity EvaGreen Mastermix (Qiagen), 8.5µL ddH2O, 0.25µL of each primers (10 µM) and 1µL of template DNA at 0.5ng/µL (see the sample description above). The qPCR program was as follows: initial denaturation at 95°C for 2min 10s, followed by 40 cycles of denaturation at 94°C for 15s, annealing at 55°C for 15s and extension at 72°C for 15s. This was followed by 1min at 72°C and held at 4°C until the amplified samples were removed from the qPCR machine. The plate with sperm and blood samples also included the standard dilutions which were used to calculate starting concentrations of the DNA samples. The change in fluorescence intensity was measured at the end of each extension step, and results were analyzed using the CFX manager software (version 3.1, Bio-Rad). Quantification cycle (Cq) values were determined by manually establishing a threshold line within the exponential amplification phase across both amplification plots.

### dPCR

Samples were analyzed on a nanoplate-based QIAcuity Digital PCR System (QIAGEN, Hilden, Germany) using the automated workflow provided by the QIAcuity Software Suite (v.2.1). dPCR reactions were performed in a 10 µL final volume consisting of 3.33 µL QIAcuity EvaGreen

Mastermix (QIAGEN, Hilden, Germany), 5.44 µL ddH2O, 0.13 µL of forward and reverse primers (10 µM) and 1 µL template DNA at 0.5ng/µL (see the sample description above). dPCR reactions were then dispensed to 96-well 8.5k QIAcuity Nanoplates (QIAGEN, Hilden, Germany), to divide all single samples into 8,500 of individual partitions before end-point PCR. The dPCR plates used in this study resulted in half as many partitions as ddPCR. Reaction conditions were as follows: 2 min initial denaturation at 95°C, followed by 35 cycles of denaturation for 15s at 95°C, 15s annealing at 55°C, and 15s extension at 72°C, with a final step of 5 min at 40°C. Results were analyzed using the QIAcuity Software Suite (v.2.1.8.23, Qiagen). Green channel was used to detect the EvaGreen® fluorophore. Exposure duration was 500ms for sperm and blood samples, and 300ms for the synthetic mtDNA fragment. The common fluorescence intensity (RFU) threshold to identify positive and negative samples was set at 57 for the analysis of sperm and blood samples, and at 30 for synthetic mtDNA analysis.

### ddPCR

Samples were analyzed on a Bio-Rad QX200 ddPCR System. ddPCR reactions were performed in a 20 µL final volume, consisting of 10µL Bio-Rad ddPCR EvaGreen supermix, 0.25 µLof forward and reverse primers (10µM), 8.5 µL of ddH2O and 1µL template DNA at 0.5ng/µL (see the sample description above). After vigorous mixing, each reaction was pipetted into a DG8 Droplet Generator Cartridge, and mixed with 70 µL of Droplet Generation Oil for EvaGreen. Droplets were then generated using the QX200 Droplet Generator (Bio-Rad), and a final 40µL volume of droplets for each sample was carefully transferred to a ddPCR 96-well plate, later sealed with pierceable foil using a PX1 PCR Plate Sealer (Bio-Rad) before end-point PCR. The specifications of the system ensure the partitioning of the sample into 20,000 droplets. PCRs were performed on a BioRad CFX96 Real-Time System (Bio-Rad Laboratories, California, United States). DdPCR conditions were as follows: 10 min at 95°C, followed by 40 cycles of denaturation for 30 s at 94°C and annealing at 55°C for 1 min, with ramp rate of 2°C/s, followed by 10 min at 98°C and a hold at 8°C. Droplets were then read on a QX200 droplet reader (Bio-Rad). Quantification data were checked using the Bio-Rad QuantaSoft software (v.1.7.4.0917). Thresholds for positive signals were determined according to QuantaSoft software instructions, and all droplets above the fluorescence threshold (10,000) were counted as positive events, those below it being counted as negative events.

### Estimation of the Limit of Detection and Limit of Quantification

The MIQE guidelines [38] define the Limit of Detection (LOD) as the lowest concentration at which 95% of the positive samples are detected. The definition of the Limit of Quantification (LOQ) varies between different authors, but is usually defined as the lowest concentration at which replicates show a CV of or less than 35% [39, 40]. To establish the LOD and the LOQ, a serial dilution was performed using the synthetic mtDNA fragment (see the primers and synthetic oligo design section). The starting point of the dilution series was 2.50E^-04^ ng/µL, followed by a 10-fold dilution series. Therefore, the serial dilution ranged from 2.50E^-04^ ng/µL to 2.50E^-12^ ng/µL, and included 9 dilution points. Ten technical replicates of each dilution point were analyzed with each quantification platform, and a minimum of 4 negative controls were included on each plate. The serial dilution further allowed us to quantify the mtDNA detected with qPCR using the standard curve generated. The LOD for ddPCR, dPCR and qPCR were calculated using dedicated R scripts as in Hunter *et al*., 2017. We also followed the method outlined by Klymus et al. [42] to assess the LOD and LOQ for qPCR assays. Thus, the LOD for qPCR was calculated using both methods [41, 42] to assess whether they will return similar results. Both scripts resulted in the same LOD for qPCR. The LOQ for ddPCR and dPCR were estimated following the threshold and methods described in [39, 40].

### Statistical analyses

In the qPCR analysis, the concentration of copy number per µL was estimated using the standard curve generated from the serial dilution of the synthetic mtDNA fragment [38]. In the dPCR and ddPCR, the copy number per µL was calculated using the QIAcuity Software Suite and QuantaSoft respectively, using the fraction of positive and negative partitions based on Poisson distribution [43]. Coefficient of Variation (later referred to as CV) was calculated for quantification results as the ratio between the standard deviation and the mean multiplied by 100. All statistical analyses were performed in R v4.3.2 using RStudio/2023.12.0+369 and the packages tidyverse v2.0.0 (Wickham *et al*., 2019), and vegan v2.6-4 [45]. The methods were compared using one-way analysis of variance (ANOVA), variance ratio test (f-test) and either Student’s t-test or Welch t-test depending on the results of the f-test. Plots were generated using ggplot2 v3.4.4 [46].

### Results

The primers developed in this study were found to be specific for passerine birds and performed well across qPCR, dPCR and ddPCR. The negative controls did not amplify with either method, except for one replicate during the analysis of the serial dilution using synthetic mtDNA to establish the LOD and LOQ on the ddPCR platform, likely due to a pipetting error. The average number of partitions generated was 8,050 with dPCR and 16,082 with ddPCR (see supplementary table S5 for all quantification information). These numbers follow the manufacturers’ specifications (see details provided in Methods section). Both dPCR and ddPCR platforms could not reliably quantify high concentration of synthetic oligo, as both platforms reached saturation due to excessive numbers of positive partitions. For this reason, we excluded high concentrations of the dilution series (i.e. dilution which led to a number of positive partitions exceeding 95%) from the downstream analysis. In contrast, qPCR was not able to detect and quantify the three lowest dilution points (Cq > 39), therefore these lower dilutions were omitted from the analysis.

### LOD and LOQ

qPCR Efficiency % was ranging from 94.5 to 116.3, slope from -3.462 to -2.984, Y intercept from 7.411 to 38.623, and R^2^ from 0.968 to 0.976 for synthetic and real mtDNA, respectively. For qPCR, the LOD was established at 2.42 copies/µL and LOQ was established at 4.9 copies/µL. dPCR had an LOD at 0.51 copies/µL and LOQ at 2.12 copies/µL, while ddPCR had an LOD at 0.63 copies/µL and LOQ at 1.23 copies/µL. We investigated the correlation between the measured concentration and the expected concentration of the dilution series (Fig. 2). Only dPCR and ddPCR were compared, as the dilution series was used later to convert qPCR Cq values into concentrations. In both methods, we found that CV decreased as the measured concentration increased, and increased when the measured concentration decreased (Table 1).

**Figure 2.**
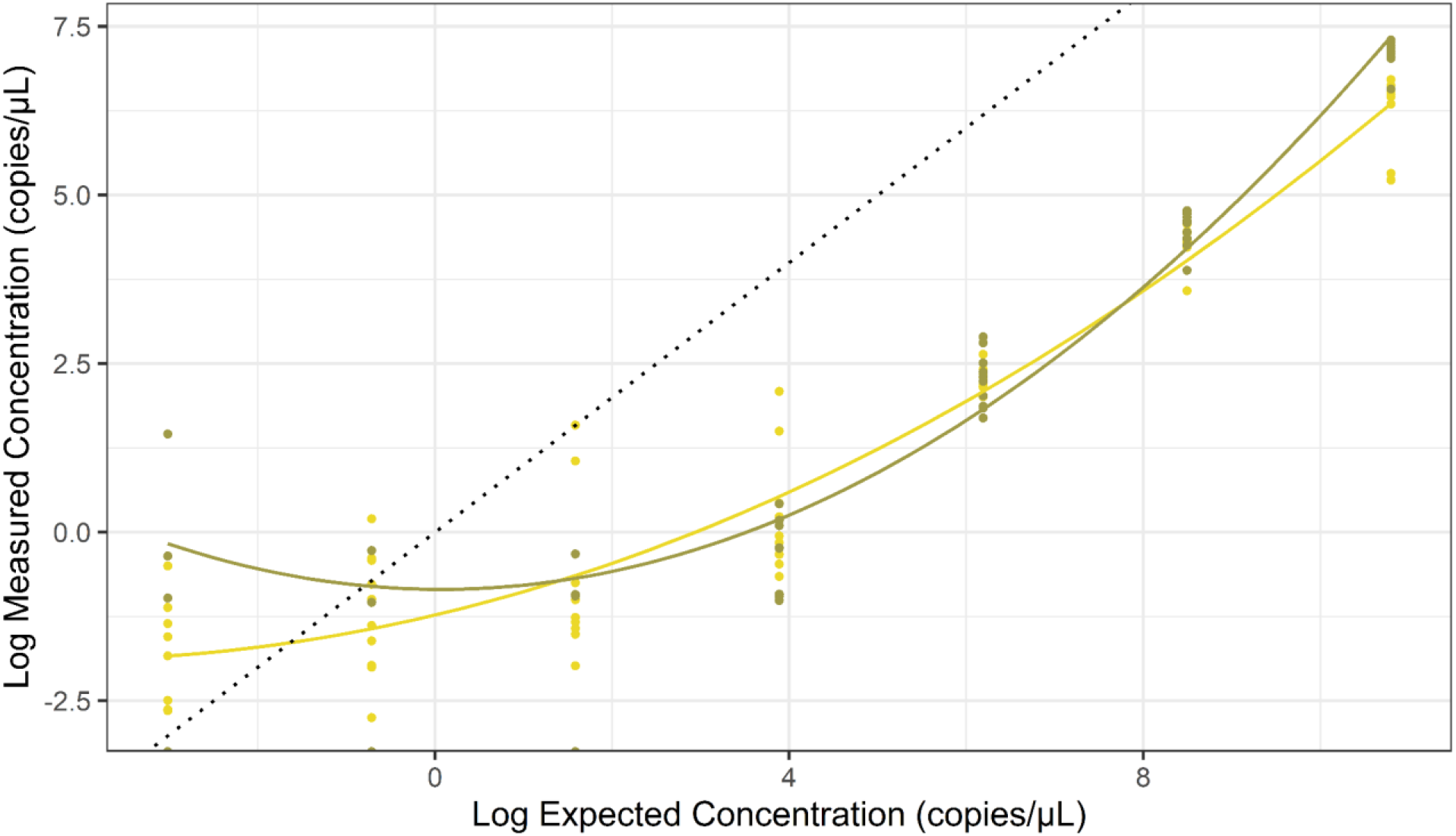
The relationship between log-transformed expected copy number of the synthetic oligo and log-transformed copy numbers obtained with dPCR (olive green) and ddPCR (yellow). Data from seven dilution points (concentration range 2.50E^-06^ to 2.50E^-12^ (ng/µL)) were used to generate this figure. The black dotted line shows where the expected concentration is equal to the observed concentration. Polynomial regression lines were fitted to the concentrations.

### Sperm and blood mtDNA

The comparison of copy numbers estimated using the three methods (Fig. 3) revealed similar mean sperm mtDNA copy number for dPCR with 310.81 copies/µL and ddPCR with 298.26 copies/µL, and not statistically different from the qPCR measurement with 507.13 copies/µL (ANOVA p>0.05).

**Figure 3.**
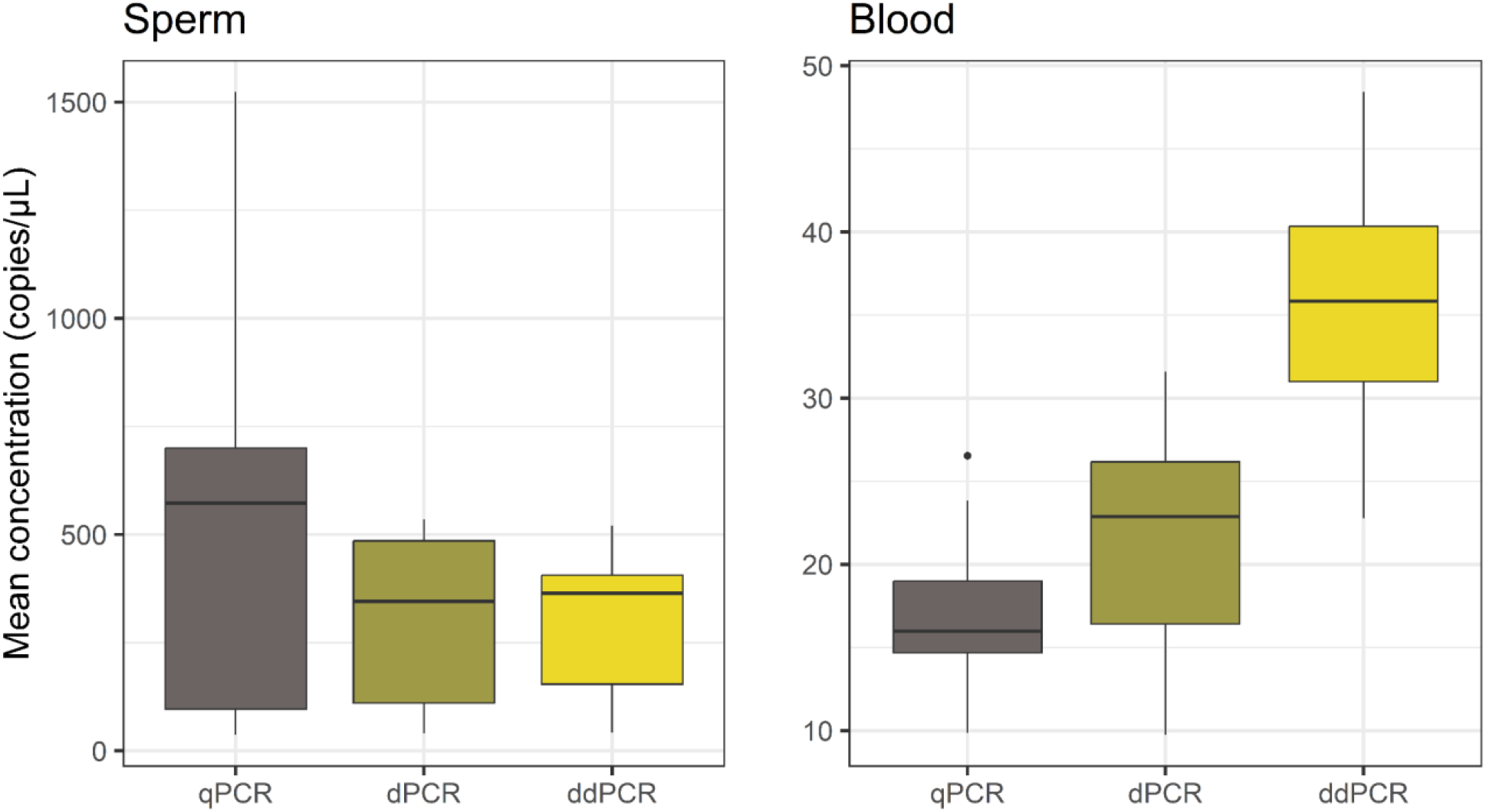
Comparison of qPCR, dPCR and ddPCR quantification of mtDNA copy number in sperm and blood samples. Each box plot represents 10 biological samples with three technical replicates each.

When comparing measured blood mtDNA concentration across the three platforms, mean concentration was 17.19 copies/µL for qPCR, 21.84 copies/µL for dPCR and 36.14 copies/µL for ddPCR. Mean ddPCR concentration was significantly different from dPCR and qPCR (ANOVA p <0.05), while the latter two showed no significant difference (ANOVA p >0.05).

The calculated copy number of samples and replicates were used to calculate CV across qPCR, dPCR and ddPCR (Fig. 4). Mean CV across methods was < 50%. For sperm samples, mean CV was 23% for qPCR, 25% for dPCR, and 9% for ddPCR. While blood samples had a higher CV with qPCR (30%) and dPCR (40%), and lower with ddPCR (7.5%). The differences in qPCR and dPCR CV were not significant (ANOVA p>0.05), while for both types of mtDNA quantification, ddPCR showed the lowest variation between replicates and was significantly different from the other two quantification methods (ANOVA p<0.05).

**Figure 4.**
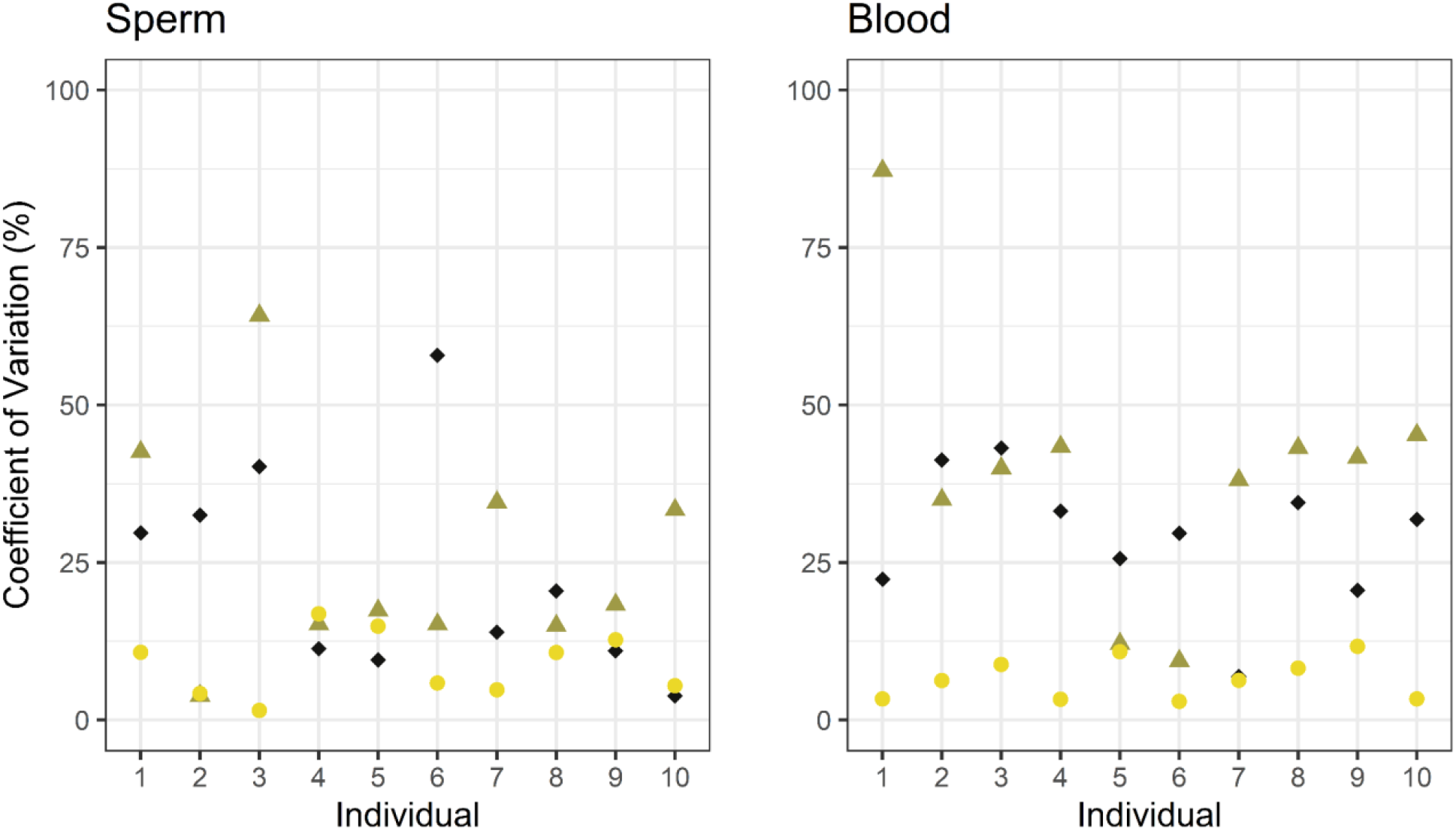
Comparison of coefficient of variation in mtDNA copy number estimation in sperm and blood across samples, every sample was quantified in triplicate. qPCR (black diamond), dPCR (olive green triangle) and ddPCR (yellow circle).

Finally, we explored the coefficient of variation of each blood and sperm samples across the three methods (Figure 5), and found that CV decreased as the concentration increased. Lower CVs were obtained when measuring sperm mtDNA samples than blood, especially for qPCR and dPCR.

**Figure 5.**
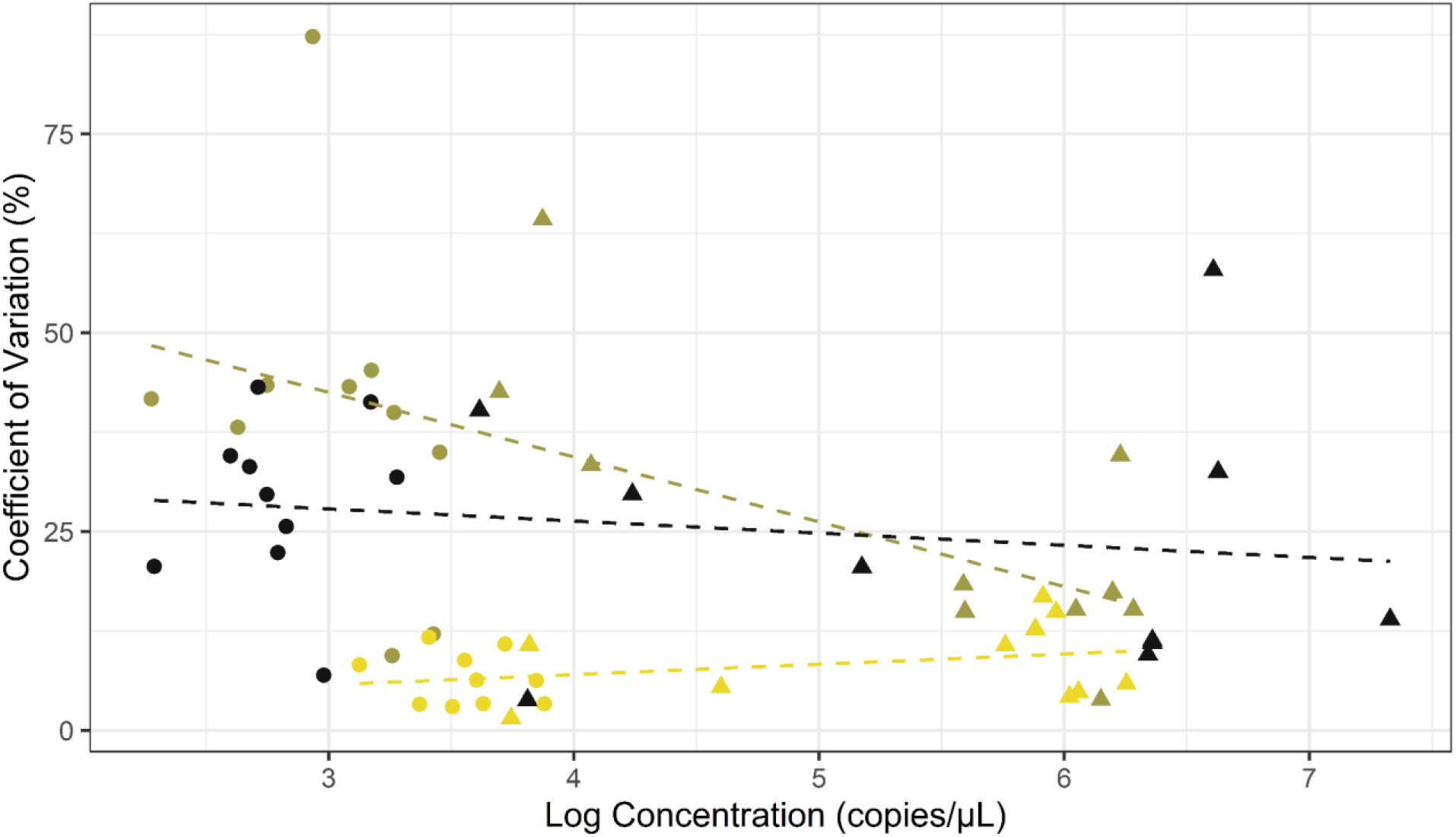
The relationship between coefficient of variation (CV) and log-transformed mean mtDNA concentration measurement of blood and sperm samples across the three quantification platforms. Regression lines for each method were added to show how CV changes with the increase of concentration • blood samples; ▴ sperm samples. Black - qPCR, olive green - dPCR, yellow - ddPCR. Overall, the lowest CV is exhibited by ddPCR.

## Discussion

In this study, we investigated the accuracy and repeatability of qPCR, dPCR and ddPCR quantification platforms to assess their reliability in quantifying mtDNA targets in avian blood and sperm samples. We first investigated the LOD and LOQ of each method using a 10-fold serial dilution of synthetic mtDNA [21]. While our LOD and LOQ measurements for qPCR, dPCR and ddPCR are in agreement with other previously published studies [39, 47], using a 5-fold dilution series with more dilution points could increase the measurements at lower concentration, therefore further improving the resolution of LOD and LOQ. Interestingly, the highest concentration of the dilution series led to saturation of both dPCR and ddPCR platforms, therefore impeding their measurements. This was not the case for qPCR, suggesting that qPCR might be more reliable when quantifying mtDNA in highly concentrated samples. However, we could not detect the target mtDNA of the three lowest concentrations from the dilution series (4.8, 0.48 and 0.048 copies/µL) using qPCR. This was not the case for both dPCR and ddPCR, suggesting that the latter two were more appropriate when quantifying low-concentration samples. This is in accordance with various studies highlighting the higher sensitivity of both dPCR and ddPCR compared to qPCR [31, 48, 49].

We found a higher copy number in sperm mtDNA compared to blood mtDNA. This can be attributed to a greater amount of mtDNA present in sperm cells compared to bird blood cells, in which mtDNA is detectable only in very low levels [6]. This accounts for the nearly tenfold difference in mtDNA concentrations we observed between the two types of cells. When comparing the three methods to measure concentrations of sperm samples that are known to have a relatively higher mtDNA copy number, we found no significant differences between platforms. mtDNA was quantified efficiently in all sperm samples even when a low amount of input DNA (0.5ng) was used in the reactions, which is important when working with wild birds with variable and often rather low volumes of ejaculates (reviewed in [50]). However, we noticed lower variation between replicates with ddPCR than with the other two methods. In contrast, blood samples have a lower mtDNA content, and even though all three methods were able to quantify the target mtDNA, we found a significant difference between quantification with ddPCR and the other two methods. This might be due to the increased reliability of ddPCR when quantifying samples with a low concentration of the target DNA, as ddPCR exhibited a much lower CV compared to qPCR and dPCR.

The similar quantification values across sperm samples suggest that all methods perform equally well to accurately and reliably measure concentration in relatively highly concentrated samples.

This corresponds to the optimal scenario (Fig. 1, scenario A), with qPCR, dPCR and ddPCR all able to accurately and repeatedly quantify the target mtDNA across replicates. However, when the level of target DNA decreases, we observe variation in quantification values between the platforms and between replicates (Fig. 3 and 5). This is leading to a less optimal scenario (Fig. 1, scenario C), where variability increases and leads to inaccurate and non-repeatable quantification across replicates. As observed with the serial dilution, when the level of target DNA decreases even more, qPCR falls into the least optimal scenario, where target detection is stochastic across replicates and quantification is incorrect, leading to unreliable results (Fig. 1, scenario D). However, it should be noted that each of the three methods described in this study has its own strengths and weaknesses, making it important for researchers to choose the most appropriate approach based on specific goals, study questions and systems. Nonetheless, qPCR requires an additional step - a reference gene or a series of standard dilutions of known concentrations, which are then used to estimate the concentration of target DNA. dPCR and ddPCR methods, on the other hand, eliminate the need for the standard curve, as the concentration values are readily available at the quantification endpoint based on Poisson statistics [28, 29]. As mentioned previously, both dPCR and ddPCR exhibited good amplification at low concentrations, and at the lowest concentration, ddPCR was more repeatable with lower CV of technical replicates. Nevertheless, it is important to consider that stochastic effects occur at low concentrations, potentially leading to the occurrence of false positive droplets. As in many laboratory processes, the human factor can be a potential limitation, and contamination or errors can happen. To mitigate this, it could be advised to employ an automated pipetting system, as it can reduce these effects, especially when working with low volume samples.

In conclusion, the choice between the three different quantification platforms investigated in our study mainly rely on methodological tradeoffs which will vary depending on the research questions. The qPCR platform has been used for decades and still is widely used across many fields of research due to its availability and broad applicability [38]. It is often more easily accessible than dPCR or ddPCR, as these more novel tools are more costly. When aiming to analyze samples showing very high levels of target DNA, qPCR quantification will be the most appropriate technique to use and will show high repeatability and accuracy, while both dPCR and ddPCR platforms could be saturated by such highly abundant targets. However, when aiming to analyze moderately high levels of target DNA, we found that the use of any of the three investigated methods will lead to accurate and repeatable results, although ddPCR shows lower variation than both qPCR and dPCR. Finally, in the case of low target quantification, we recommend the use of either dPCR or ddPCR, as both platforms show reliable and similar quantification results, with ddPCR again showing significantly lower variation than dPCR.

## Acknowledgments

We would like to thank Birgitte Lisbeth Graae Thorbek, Audun Schrøder-Nielsen and Jarl Andreas Anmarkrud from the DNA lab at the Natural History Museum at the University of Oslo for their support and assistance in the lab. We also thank SeAnalytics AB for processing the dPCR samples.

## Author Contribution

Sampling: EHL, JTL; Laboratory analysis: LB and QM; Data curation: LB and QM ; Statistical analysis: LB and QM; Original draft: LB and QM; Review and editing: LB, JTL, AJ, EHL and QM; Funding acquisition: JTL; Supervision: JTL, AJ, EHL and QM.

## Data Availability Statement

Raw data can be found in the supplementary information section.

