## Supplementary information for "Assessing reliability and accuracy of qPCR, dPCR and ddPCR for estimating mtDNA copy number in songbird blood and sperm cells"

**Table S1.** Sperm and blood samples used in the study. Ten sperm samples from 8 individuals and 10 blood samples from 10 individuals were used in the study. Two individuals were sampled twice for sperm (NHMO-BI-108302 and NHMO-BI-108464).

| Sample type | Accession number | Species | Collection date | Collection location |
| --- | --- | --- | --- | --- |
| Sperm | NHMO-BI-108302 | *Spinus spinus* | 02/06/2022 | 61.226 N \| 9.040 E |
| Sperm | NHMO-BI-108302 | *Spinus spinus* | 02/06/2022 | 61.226 N \| 9.040 E |
| Sperm | NHMO-BI-108301 | *Spinus spinus* | 02/06/2022 | 61.226 N \| 9.040 E |
| Sperm | NHMO-BI-108479/2-S | *Spinus spinus* | 13/06/2022 | 61.226 N \| 9.040 E |
| Sperm | NHMO-BI-108464/1-S | *Spinus spinus* | 02/06/2022 | 61.226 N \| 9.040 E |
| Sperm | NHMO-BI-108464 | *Spinus spinus* | 02/06/2022 | 61.226 N \| 9.040 E |
| Sperm | NHMO-BI-106779/1-S | *Spinus spinus* | 11/06/2021 | 61.226 N \| 9.040 E |
| Sperm | NHMO-BI-106744/1-S | *Spinus spinus* | 11/05/2021 | 61.226 N \| 9.040 E |
| Sperm | NHMO-BI-106748/1-S | *Spinus spinus* | 17/05/2021 | 61.226 N \| 9.040 E |
| Sperm | NHMO-BI-106746/1-S | *Spinus spinus* | 13/05/2021 | 59.744 N \| 10.820 E |
| Blood | NHMO-BI-106927/1-B | *Spinus spinus* | 13/05/2021 | 59.744 N \| 10.820 E |
| Blood | NHMO-BI-108472/1-B | *Spinus spinus* | 13/06/2022 | 61.226 N \| 9.040 E |
| Blood | NHMO-BI-108479/1-B | *Spinus spinus* | 13/06/2022 | 61.226 N \| 9.040 E |
| Blood | NHMO-BI-108493/1-B | *Spinus spinus* | 22/06/2022 | 61.226 N \| 9.040 E |
| Blood | NHMO-BI-109025/1-B | *Spinus spinus* | 13/07/2022 | 61.226 N \| 9.040 E |
| Blood | NHMO-BI-109030/1-B | *Spinus spinus* | 13/07/2022 | 61.226 N \| 9.040 E |
| Blood | NHMO-BI-109034/1-B | *Spinus spinus* | 13/07/2022 | 61.226 N \| 9.040 E |
| Blood | NHMO-BI-109036/1-B | *Spinus spinus* | 13/07/2022 | 61.226 N \| 9.040 E |
| Blood | NHMO-BI-109039/1-B | *Spinus spinus* | 13/07/2022 | 61.226 N \| 9.040 E |
| Blood | NHMO-BI-109040/1-B | *Spinus spinus* | 13/07/2022 | 61.226 N \| 9.040 E |

**Table S2.** GenBank accession numbers of the sequences used for primer design.

| **Species** | **GenBank accession numbers** |
| --- | --- |
| Hawfinch (*Coccothraustes coccothraustes*) | KX109614.1; NC_025614.1; KM078789.1 |
| Great tit (*Parus major*) | HQ417139.1; FJ465247.1; AJ224494.1; KP137624.1; NC_040875.1 |
| Common reed bunting (*Emberiza schoeniclus*) | KP877661.1; OQ274893.1; MW417350.1; NC_062087.1; MT081419.1 |
| Eurasian chaffinch (*Fringilla coelebs*) | HQ284765.1; NC_025599.1; MN122830.1; KM078769.1 |
| Eurasian bullfinch (*Pyrrhula pyrrhula*) | HQ284732.1; NC_025625.1; KM078804.1; HQ284731.1; HQ284730.1 |
| Bluethroat (*Luscinia svecica*) | MN122892.1 |
| Eurasian siskin (*Spinus spinus*) | NC_015198.1; HQ915866.1 |

**Table S3.** Sperm samples used for PCR and ddPCR validation of the primers.

| **Species** | **Accession number** | **Collection date** | **Collection location** |
| --- | --- | --- | --- |
| *Spinus spinus* | NHMO-BI-106957  NHMO-BI-108466  NHMO-BI-108463 | 28/04/2021  02/06/2022  02/06/2022 | 59.744 N \| 10.820 E  61.226 N \| 9.040 E  61.226 N \| 9.040 E |
| *Turdus pilaris* | NHMO-BI-106959 | 08/06/2021 | 61.25 N \| 9.06 E |
| *Phylloscopus trochilus* | NHMO-BI-106966 | 17/06/2021 | 61.22 N \| 9.04 E |
| *Luscinia svecica* | NHMO-BI-100507 | 24/05/2018 | 61 25 N \| 8 52 E |
| *Fringilla montifringilla* | NHMO-BI-106797  NHMO-BI-108304 | 21/05/2021  04/06/2022 | 61.226 N \| 9.040 E  61.226 N \| 9.040 E |
| *Pyrrhula pyrrhula* | NHMO-BI-104533 | 19/06/2020 | 61.226 N \| 9.040 E |
| *Regulus regulus* | NHMO-BI-108611  NHMO-BI-108615 | 07/05/2022  13/05/2022 | 59.98 N \| 10.74 E  59.99 N \| 10.79 E |
| *Coccothraustes coccothraustes* | NHMO-BI-107668  NHMO-BI-107669 | 19/05/2022  19/05/2022 | 59.744 N \| 10.820 E  59.744 N \| 10.820 E |
| *Emberiza schoeniclus* | NHMO-BI-108669  NHMO-BI-108534 | 21/05/2022  15/05/2022 | 61 25 N \| 8 52 E  61.226 N \| 9.040 E |

**Table S4.** The LOD and LOQ values for synthetic DNA concentrations with the three different PCR methods. The method outlined in [1] was used to assess LOD and LOQ for qPCR. The LOD for ddPCR, dPCR and qPCR has been calculated using the script provided in the supplementary material as in [2]. Both methods show  similar LOD values for qPCR: 2.4 copies/µL using the method described in [1] and 2.417 copies/µL using the method described by Hunter et al. [2]. To establish  LOQs for dPCR or ddPCR, we followed the method outlined in [3] and [4], and considered LOQ as the lowest concentration at which the CV is ≤ 35%.

| **Method** | **LOD (copies/µL)** |  | **LOQ (copies/ µL)** |
| --- | --- | --- | --- |
| dPCR | 0.51 |  | 2.12 |
| ddPCR | 0.63 |  | 1.23 |
| qPCR | 2.42 |  | 4.9 |

**Table S5.** Calculated copy number of synthetic mtDNA. Based on the molecular weight (30854 g/mole) of the synthetic oligo, the number of copies (molecules) for the different dilutions of the synthetic DNA was estimated using the formula provided below the table.

| **Dilution steps (ng/µL)** | **Copy number (copies/µL)** |
| --- | --- |
| 0.00025 | 4879513 |
| 0.000025 | 487951.3 |
| 2.5E-06 | 48795.13 |
| 2.5E-07 | 4879.513 |
| 2.5E-08 | 487.9513 |
| 2.5E-09 | 48.79513 |
| 2.5E-10 | 4.879513 |
| 2.5E-11 | 0.487951 |
| 2.5E-12 | 0.048795 |

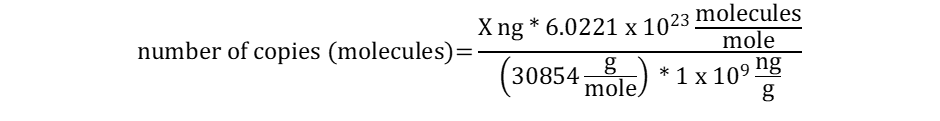

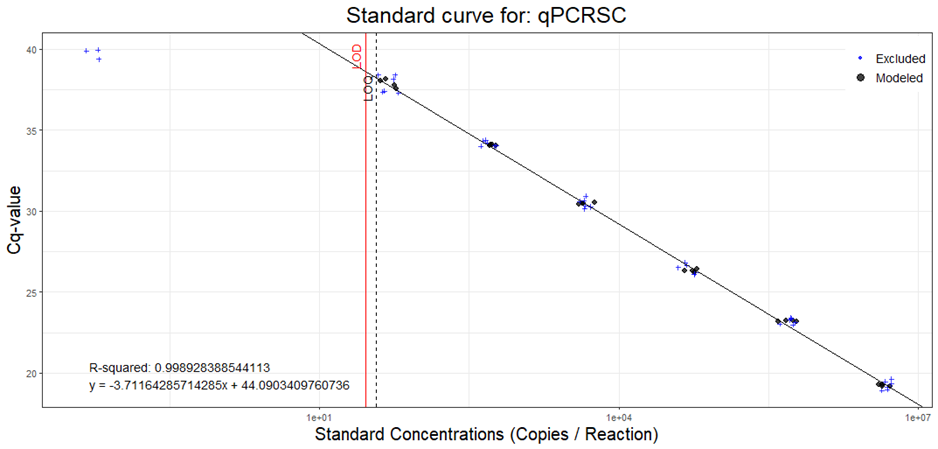

**Figure S1.** Estimated Limit of Detection (LOD) and Limit of Quantification (LOQ) for qPCR. The calculations and the figure were generated following the R script provided in [1].

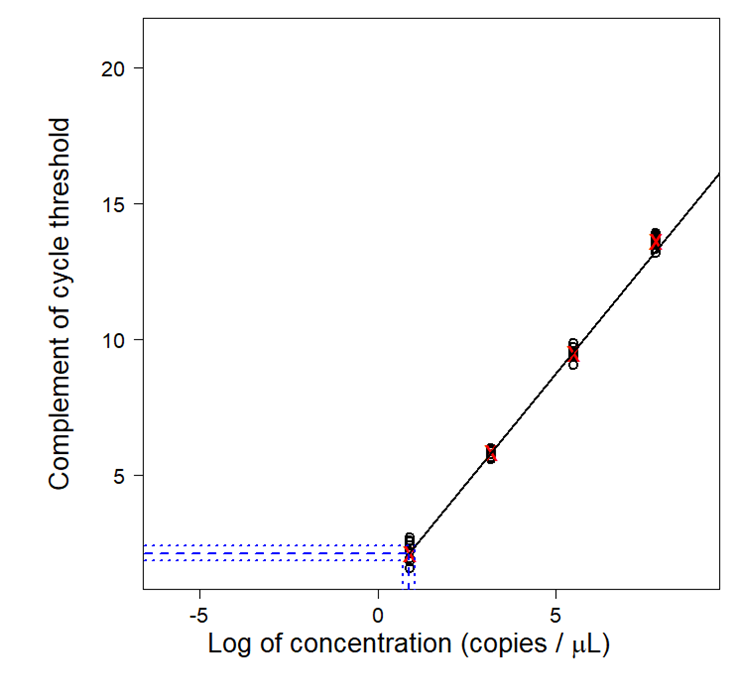

**Figure S2.** Limit of Detection estimation for qPCR. Figure and calculations were performed  following the method outlined in [2].

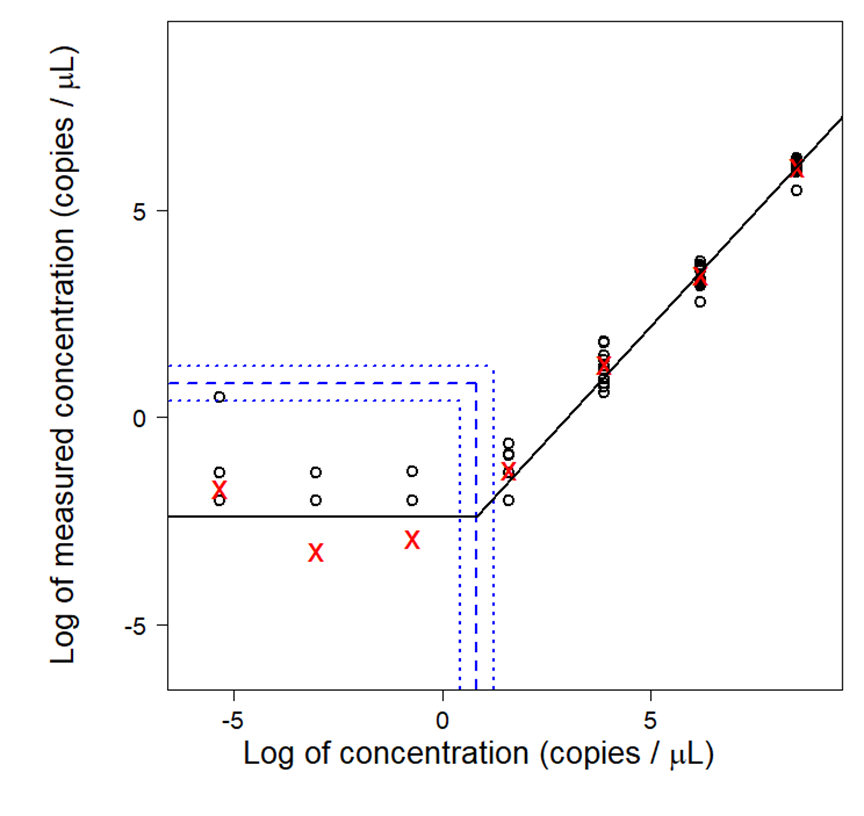

**Figure S3.** Limit of detection estimation for dPCR. Figure and calculations were performed following the method outlined in [2].

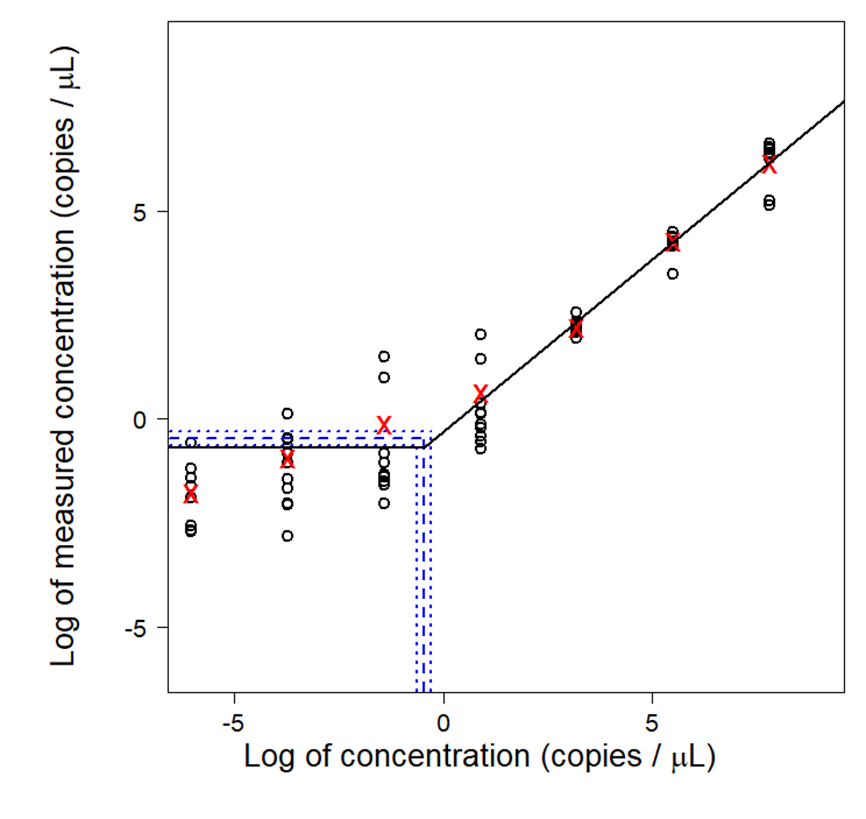

**Figure S4.** Limit of Detection estimation for ddPCR. Figure and calculations were performed following the method outlined in [2].

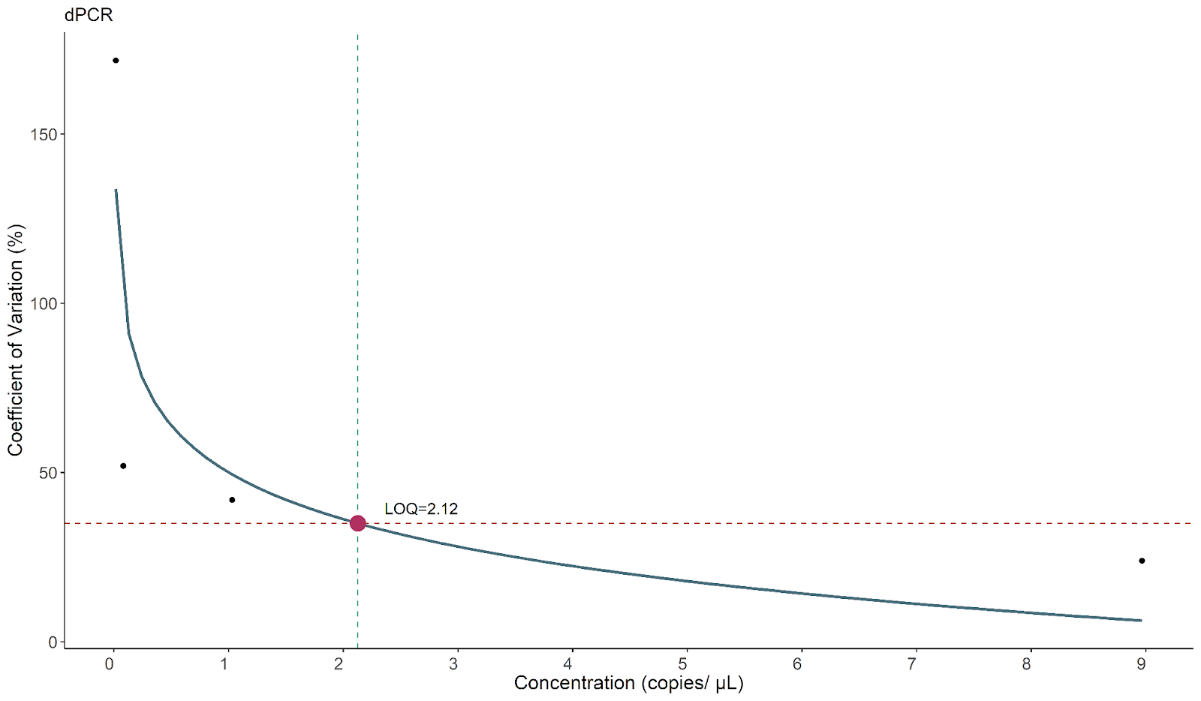

**Figure S5.** Limit of Quantification (LOQ) of dPCR.

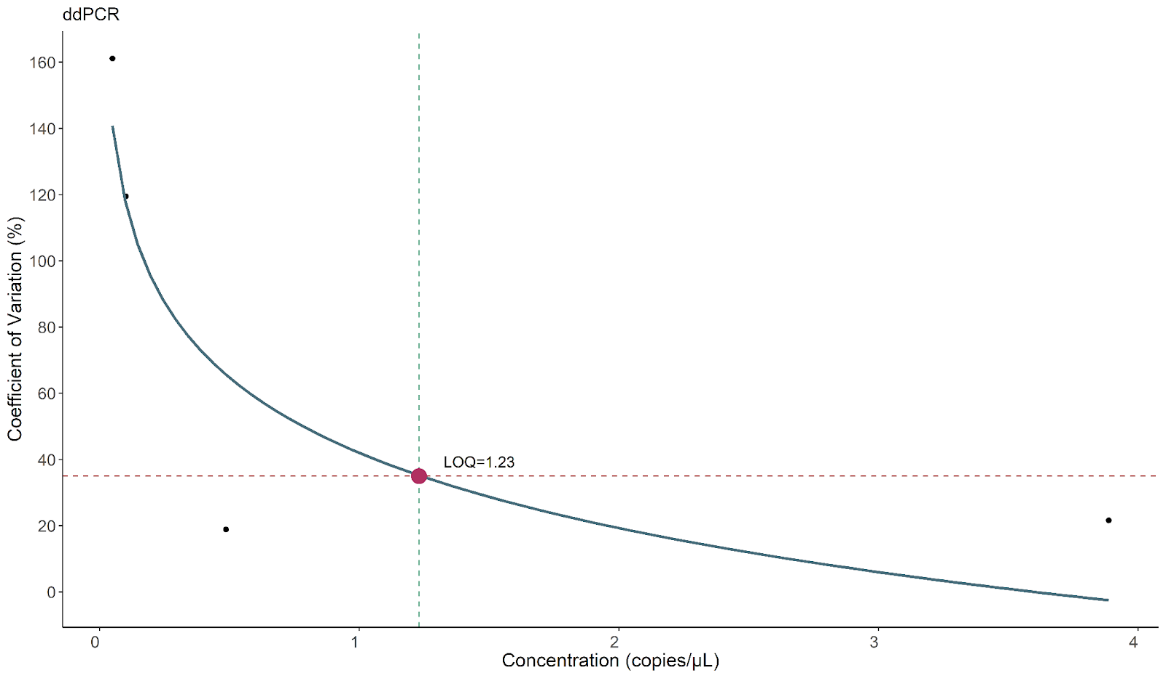

**Figure S6.** Limit of Quantification (LOQ) of ddPCR.
